# Context-Dependent Flux Coupling via Conserved Small-Molecule Regulatory Structures

**DOI:** 10.1101/2021.08.27.458000

**Authors:** Christian Euler, Radhakrishnan Mahadevan

## Abstract

Small-molecule regulation modulates enzyme activity and is widespread in metabolic networks. However, the organization of small-molecule regulatory networks and its generalized role is not well understood. We analyze the structure of the genome-wide *Escherichia coli* small-molecule regulatory network (SMRN) to reveal that it optimizes controllability in the metabolic network. This is achieved by conserved, highly overabundant incoherent feedforward loops. Using multi-omics data, we characterize loop examples in central carbon metabolism. These use signals from hypothesized flux-sensing metabolites phosphoenolpyruvate, *α*-ketoglutarate, citrate, and malate to distinguish between glycolysis, gluconeogenesis, and glyoxylate shunt activity to differentially couple fluxes across these major modes of metabolism. Our results suggest that coupling of fluxes by direct modulation of enzyme activity is an emergent property of the SMRN that depends heavily on both regulatory structure and metabolic context via the metabolome, and further that flux sensing and coupling may be a global property of the metabolic network.

## Introduction

Small-molecule regulation (SMR) remains relatively unexplained at the systems level [19]. Detailed mechanistic and structural explanations of the major forms of SMR – allostery and the various modes of enzyme inhibition – exist and have been validated for individual proteins in numerous biological systems [17, 29]. However, unlike other metabolic regulatory mechanisms, the underlying design principles guiding SMR organization into local and global structures have yet to be explored. For example, structure-function analyses have been performed on transcriptional regulatory networks (TRN) at local and global scales, and these have enabled both the identification of structural motifs [25] and the mechanisms by which they control cellular behaviour [12]. Our aim is therefore to close the SMR knowledge gap by elucidating how its topological organization within metabolism generally influences control of metabolic fluxes.

The first small-molecule regulatory network (SMRN) reconstruction demonstrated that metabolites with similar structures affect the same groups of enzymes, a fact that has recently been recapitulated experimentally [13, 33]. A more recent reconstruction has revealed that metabolic branchpoints are *not* more likely than any other local structure to be regulated by SMR, but that biosynthetically expensive metabolites are often effectors and their reactions are often regulated [34]. Experimental work has also shown that SMR may be far more ubiquitous than is currently suspected because there are significant barriers to characterizing the phenomenon *in vivo*. Indeed, several approaches to evaluating relevant *in vivo* SMR interactions have been developed in recent years with varied success [6, 18, 20, 30, 31].

The underlying motivation for understanding systems-level SMR derives from the fact that it plays a significant role in the coordination of metabolism [3]. For example, in several organisms, transcriptional regulation cannot sufficiently explain observed metabolic responses across varied static environments [4], or responses to environmental perturbations [8], and real flux distributions may be best predicted with models that explicitly integrate SMR interactions [14]. The addition of these to kinetic models can increase predicted network stability [11], and may be sufficient to both predict glucose uptake into metabolism [27] and explain growth optimality [10]. However, the inherent nonlinearity and speed of the phenomenon makes it challenging to integrate at scale [23]. These limitations have also made analyzing the role of SMR in controlling fluxes through reaction cycles (e.g. the TCA cycle) difficult: to date, no work has explicitly addressed this. A high-level topological understanding could address these challenges by highlighting the role of SMR structures and the context in which they are expected to be relevant.

To this end, we show here that the *E coli* SMRN is dominated by novel feedforward loops that connect highly connected reactions to weakly connected reactions in the metabolic network. We develop a general steady-state model of loop function and use reference metabolomic and fluxomic datasets to show that these structures couple fluxes in specific metabolic contexts, and that this emergent flux coupling in central carbon metabolism, in particular, the TCA cycle, can distinguish between the major metabolic modes of glycolysis, gluconeogenesis, and the glyoxylate shunt to provide distinct, context-dependent control in each of these.

## Results

### Network Reconstruction

To reconstruct the *E coli* SMRN, we extracted regulator-enzyme pairs from both BRENDA and the Allosteric Database [38], then cross-referenced regulators against the metabolites in the genome-scale metabolic model (GSM) iJO1366 using their KEGG IDs to exclude any pairs containing non-native regulators. We associated the remaining pairs to reactions in the GSM using the EC number, Uniprot accession numbers, and gene rules for each reaction. Reactions and metabolites associated with the regulatory network were protected in a standard loss-free network minimization [5] to produce a bipartite network containing 955 reactions. Reactions in this network are connected to their products via directed catalytic edges, and metabolites to the reactions that consume them via directed catalytic edges (Fig 1A, black arrows). Similarly, metabolites are connected to reactions they regulate via directed regulatory edges of one of two types, inhibition (Fig 1B, red edges) or activation (Fig 1B, green arrows). Thus, this process generated two subnetworks which we call the catalytic or metabolic network and small-molecule regulatory network (SMRN), respectively (Fig 1B).

**Figure 1.**
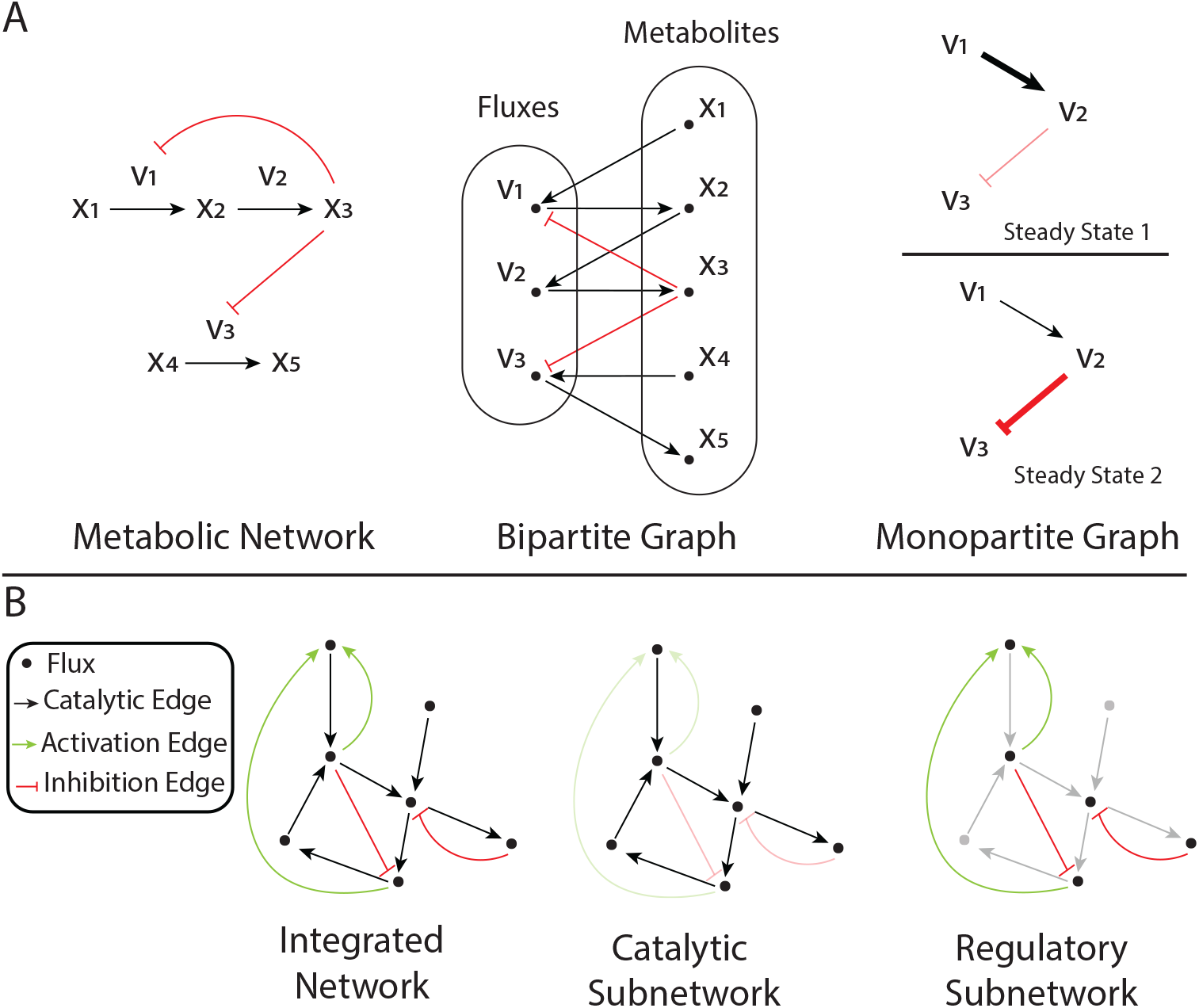
Network reconstruction and randomization. (A) Reconstruction as a monopartite sensitivity netowrk. The metabolic network is made up of fluxes, metabolites, and regulatory interactions. Since all fluxes must be connected via metabolites, this naturally yields a bipartite graph describing sensitivity interactions. For example, a positive perturbation in *x*_1_ increases *v*_1_, which in turn causes a positive perturbation in *x*_2_, etc. Conversion to a monopartite graph requires embedding information about metabolite concentrations in edges so that they have different weights (representing different interaction strengths) in different steady states. (B) The resulting integrated monopartite graph has two subnetworks: the catalytic subnetwork, equivalent to the metabolic network that is the typical unit of analysis, and the regulatory subnetwork, which is made up of only regulatory interactions.

Since we were concerned with functional coupling between reaction rates or fluxes, we converted the bipartite graphs representing each subnetwork into monopartite graphs by directly connecting reactions with a catalytic edge when the product of one reaction was the substrate of another, and with a regulatory edge when the product of a reaction was the regulator of another (Fig 1B). This novel reconstruction approach generated a sensitivity network in which nodes are rates and edges represent possible coupling relationships between them. The weights of these edges, representing coupling strength, are dependent on the steady-state concentrations of the metabolite(s) mediating coupling (Fig 1B). At a given steady-state, a perturbation to a node in the network can be propagated to other nodes to which it is connected in a concentration-dependent manner.

The resulting integrated network contains 4400 catalytic edges connecting the 955 reactions remaining after additional pruning (see Materials and Methods). Of these reactions, 623 are in the regulatory subnetwork and are connected by 3985 edges. Approximately 80% of all regulatory edges represent negative interactions, which is consistent with previous SMRN reconstructions [13, 34].

### Metabolic Interactions & Connectivity Predicts Regulatory Role in the SMRN

It is currently not understood which reactions and metabolites are favoured as small molecule regulators and which are favoured as targets of these interactions. We hypothesized that the topological features of reactions could be predictive of their regulatory roles, since topology is known to influence controllability [22]. As is commonly done to find non-random network structural features, we generated an ensemble of 1000 randomized regulatory networks using an edge-swapping method [26] to uncover such topological features and associate them to regulatory role.

We made use of topological overlap, *O*_*t*_(*i, j*), a normalized measure of shared neighbours between any two nodes, *i* and *j*, in a network, as a simplified characterization of metabolic network topology. *O*_*t*_(*i, j*) = 1 represents the case in which either all connections between *i* and *j* are shared or the connections of one is a subset of the other. In contrast, *O*_*t*_(*i, j*) = 0 represents no shared connections. Since *O*_*t*_(*i, j*) is calculated for each possible pairing of nodes, each individual node has a distribution of *O*_*t*_(*i, j*) values describing how it is connected to the network at large, and this can be used as a topological signature. An individual node with a high average *O*_*t*_(*i, j*) value, 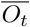, shares neighbours with a large fraction of the network, so it is more strongly connected than a node with low 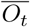. Since our network is a flux sensitivity reconstruction, strongly interacting nodes are fluxes which are influenced by and/or which influence many other fluxes.

The distribution of 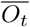 values for the catalytic network is bimodal, indicating that there are two groups of nodes in this subnetwork: a few nodes overlap more significantly with the network at large, while most nodes have little overlap with the network at large (Fig 2A). This is reflected in the topological overlap matrix (TOM) for the catalytic subnetwork, in which nodes are clustered by their *O*_*t*_(*i, j*) values: approximately one third of the network overlaps strongly, yielding a hierarchically clustered region, while two thirds of all nodes have low overlap with the rest of the network (Fig 2D, left panel). Thus, a small number of fluxes are strongly connected in the metabolic network, while most are only weakly connected, recapitulating previous characterizations of metabolic network structure [24].

**Figure 2.**
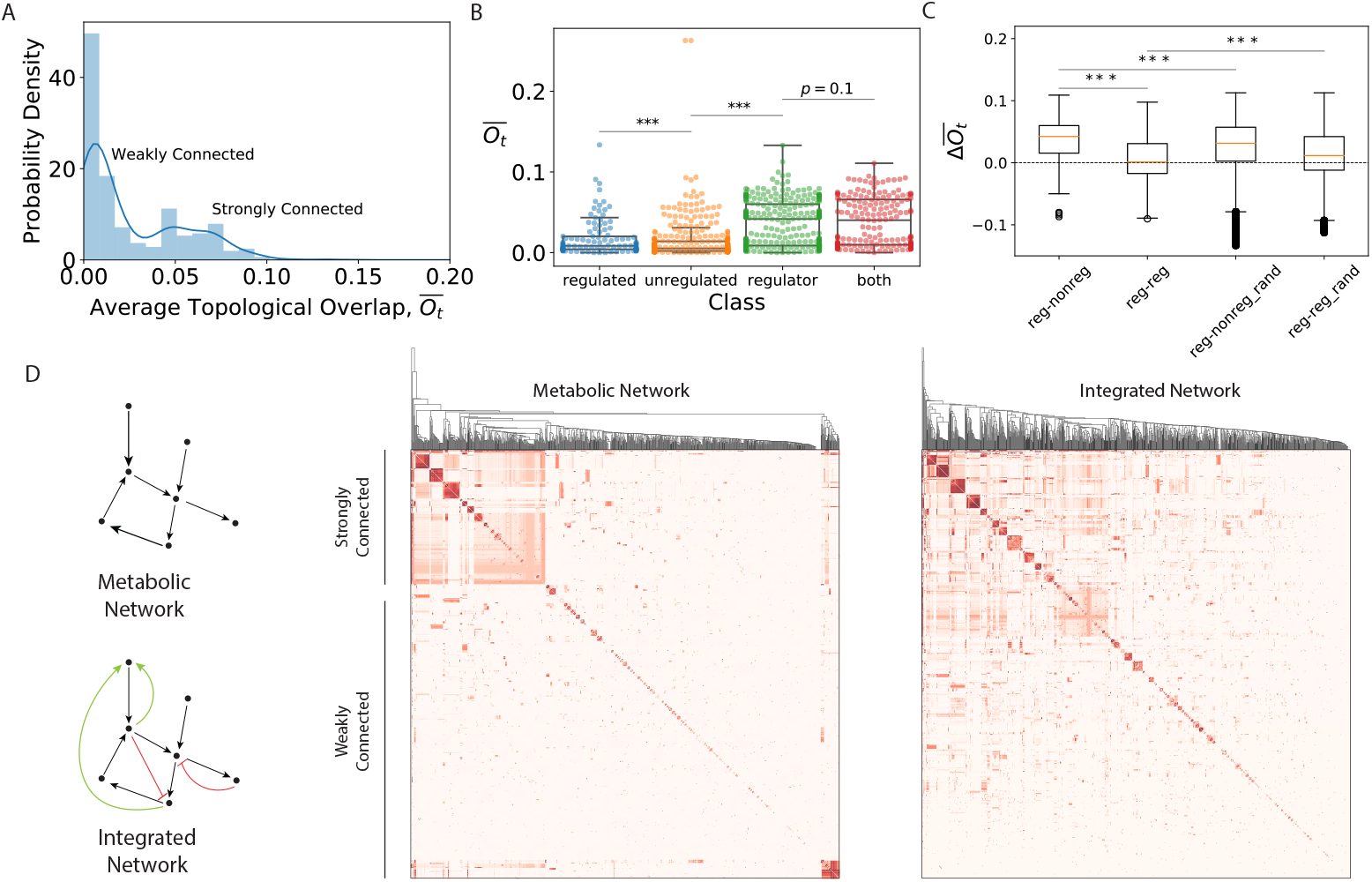
Regulatory interactions connect weakly connected nodes to strongly connected nodes in the metabolic network. (A) Average topological overlap 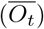 distribution for nodes in the metabolic network. Approximately one-third of all nodes are strongly connected in the network, indicated by high 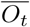, while two thirds have low 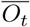. (B) 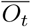 distribution by class. Nodes are regulated if they have only regulatory in-edges, unregulated if they have no regulatory edges, regulators if they have only regulatory out-edges, and both if they have regulatory in-and out-edges. Regulated nodes have lower 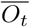 on average than unregulated nodes (Mann-Whitney U test, *p <* 0.001), and unregulated nodes have lower 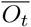 on average than regulators (Mann-Whitney U test, *p <* 0.001). Nodes which have regulatory out-edges are topologically indistinguishable whether they have in-edges as well or not (K-S test, *p* = 0.1). (C) 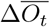 by pairing type. Regulatory-nonregulatory pairings have high 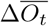 relative to regulatory-regulatory pairings (Mann-Whitney U Test, *p <* 0.001), and both of these kinds of pairings are significantly different from the same pairings predicted in the randomized networks (suffix “rand”, Mann-Whitney U test, *p <* 0.001). The regulatory network favours connections between strongly connected nodes and weakly connected nodes more often than is predicted by random chance. (D) Topological overlap matrices (TOM) for the metabolic and integrated networks. The metabolic network is hierarchically clustered, with a highly interdependent core, and many weakly connected nodes. Addition of regulatory edges reduces this hierarchy and yields significantly more between-cluster overlap.

To understand if topological connectivity predicts regulatory role, we classified nodes as ‘regulators’ if they have regulatory out-edges only, ‘regulated’ if they have regulatory in-edges only, ‘both’ if they have in-and out-edges, and ‘unregulated’ if they have no regulatory edges, and compared the distributions of 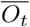 values for nodes in each class in Fig 2B. We found that ‘regulated’ nodes have the lowest 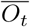 values, ‘unregulated’ nodes have slightly higher 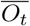 values (Mann-Whitney U Test, *p* = 5.0 *·* 10^−4^), and nodes with regulatory out-edges (‘regulator’, ‘both’) have significantly higher 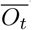 values than ‘unregulated’ nodes (Mann-Whitney U Test comparing ‘regulator’ nodes to ‘unregulated’ nodes, *p* = 3.2 *·* 10^−35^), but 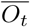 distributions which are indistinguishable from each other (K-S test, *p* = 0.17). Nodes with regulatory out-edges – i.e. reactions the produce effectors – are among the strongly connected third of all nodes in the metabolic network. This suggests that addition of regulatory interactions to the catalytic network is non-random, and likely predicted by how connected the nodes are in the metabolic network. Highly-connected nodes are sources of control signals in the SMRN, and the most weakly connected nodes are more often targets of these signals.

To highlight this finding, we calculated the difference in 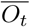 values, 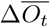, for each pairing in the SMRN, then sorted these by class in Fig 2C. 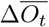 for regulatory-regulated node pairings are on average higher (0.042 ± 0.029) than those for regulated-regulatory node pairings (0.001 ± 0.03) (Mann-Whitney U Test, *p* = 1.2 *·* 10^−160^). Both types of pairings have 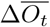 values which are significantly different (higher, lower, respectively) from those predicted by randomized ensembles (*p <* 0.001, see Materials and Methods). Thus, regulatory interactions are significantly more likely to connect nodes which are strongly connected in the metabolic network to nodes which are weakly connected in the metabolic network than they are to connect strongly connected nodes to other strongly connected nodes, and this has not likely occurred due to random chance alone. This is reflected in the TOM of the integrated network resulting from addition of the regulatory edges to the catalytic subnetwork (Fig 2D, right panel). Addition of these edges reduces the hierarchical structure of the catalytic subnetwork and increases cross-network overlap. Hence, the sources of regulatory signals are concentrated among a few highly connected nodes whereas the targets of the signals they produce are more often weakly connected nodes.

Weakly-connected nodes in the metabolic network are not coupled to many local fluxes, thus they are less likely to be controllable through metabolic interactions alone [22]. The SMRN is structured to address this problem by linking these reactions to highly-connected nodes via small-molecule regulation instead, and this appears to be a global principle of SMRN organization. Though the *de facto* understanding of SMR is as a within-pathway phenomenon, and canonically limited to simple linear feedback and feedforward regulation, our results suggest that there is in fact non-random global organization in the *E coli* SMRN, and that this structure serves to redistribute sensitivity away from the natural hierarchy that exists in the metabolic network. This renders topologically disconnected fluxes more dependent on fluxes elsewhere in the network, and suggests that globally the SMRN has been optimized to maximize controllability around structural constraints inherent to the metabolic network.

### Novel, Topologically Unique Feedforward Loops Redistribute Sensitivity

We thus wanted to understand how local structures in the SMRN distribute control in the metabolic network. Feedforward loops (FFLs) are minimal functional structures in regulatory networks [25], so we enumerated and classified the loops in the *E coli* SMRN and each network in our randomized ensemble to understand whether such loops are relevant. We found that 51.5% of all reactions in the real network are involved in the 4569 FFLs we found (Fig 3B, dashed line), while in randomized networks only 49.0 ± 0.6% of all reactions are involved in such structures (Fig 3B). The probability of this occurring by random chance alone is 1.3 ·10^−5^. FFLs are thus significantly overabundant in the *E coli* SMRN.

**Figure 3.**
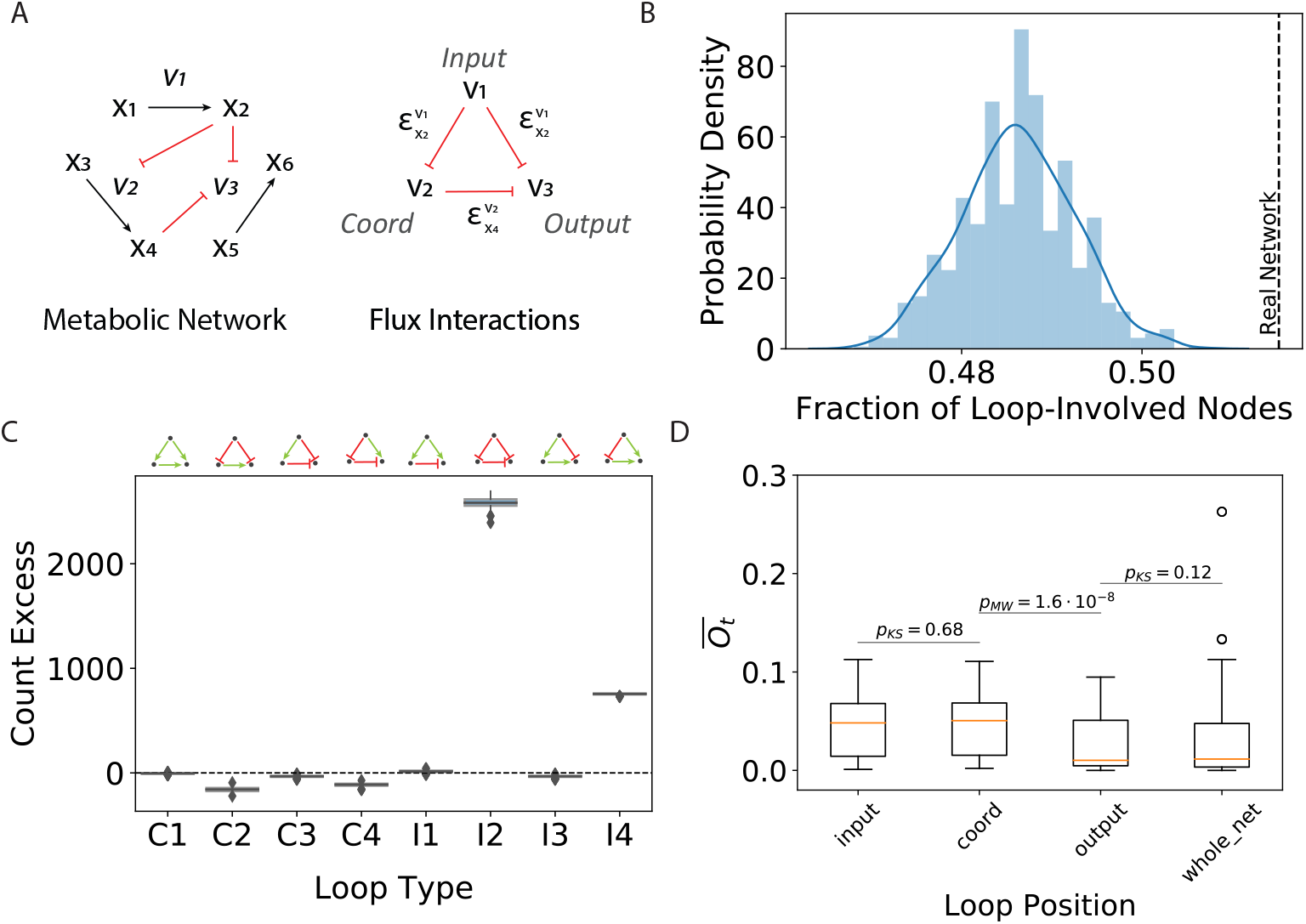
Feedforward loops (FFL) are abundant in the SMRN and have unique topological signatures. (A) FFL schematic for loops in the SMRN. The input flux produces an effector of both the coordinating and output fluxes. The coordinating flux also produces an effector of the output flux. (B) Fraction of loop-involved nodes predicted by the randomized ensemble of networks. Across 1000 randomized networks, 49.0 ± 0.6% of all reactions are involved in FFLs. In the real network (dashed line), 51.5% of all nodes are involved in FFLs. FFLs are significantly more abundant in the real network than would be predicted by random chance alone. (C) Excess loop count by type relative to randomized ensemble. The real network has significantly more I2 and I4 loops than is predicted by the randomized ensemble, indicating that these structures are resistant to negative selection and/or offer beneficial function to the metabolic network. In contrast, C2 and C4 loops are underrepresented relative to their abundance among the ensemble networks, suggesting that they are detrimental to the metabolic network and therefore do not persist over time. ‘C’ indicates coherent and ‘I’ indicates incoherent. Each loop structure is provided above the plot. (D) FFL positions are related to topological properties. Nodes at the input and coordinating positions have indistinguishable 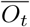 distributions (K-S test, *p* = 0.68) which are significantly higher on average than those of nodes at the output position (Mann-Whitney U test, *p* = 1.6 · 10^−8^). Output nodes are topologically indistinguishable from the network at large (K-S test, *p* = 0.12). Input and coordinating nodes are more often those which are strongly connected in the network, while output nodes are not topologically unique.

Further, we found 3514 incoherent type 2 (I2) loops and 797 incoherent type 4 (I4) loops in the real network, representing excesses of ∼ 2700 and ∼ 750, respectively, relative to those enumerated in the randomized ensembles. Based on the distributions of randomized loop counts, the probability of these excesses occurring by random chance alone is immeasurably small. Conversely, we found statistically significant deficits of ∼200 coherent 2 (C2) and and ∼ 100 coherent 4 (C4) loops in comparison to the randomized ensemble (Fig 3C, Fig S2). The over-and underrepresentation of loops relative to random chance indicates that certain local structures offer functional advantages, while others may be detrimental to metabolic control.

As in Fig 3A, Each FFL is made up of an input node, a coordinating node, and an output node. We examined the 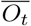 distributions for nodes in each of these positions in all of the loops to understand how they are positioned within the metabolic network. This revealed that the distribution of 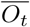 values for input and coordinating nodes are indistinguishable (K-S test, *p* = 0.68), and significantly higher than those of output nodes (Mann-Whitney U Test comparing coordinating nodes to output nodes, *p* = 1.6 *·* 10^−8^), while the 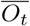 distribution for output nodes is indistinguishable from that of all nodes in the SMRN (K-S test, *p* = 0.12)(Fig 3D). Nodes at the input and coordinating positions of FFLs are those which are most strongly connected in the network, while the output nodes are not topologically distinct in the metabolic network at large. Thus, FFLs are well structured to receive and coordinate flux signals within the highly interdependent core of metabolism, and these signals control fluxes across the network without structural bias. FFL structures appear to play a significant role in redistributing control across the metabolic network, and are thus likely functional units in the SMRN.

### Feedforward Loops Differentially Couple Fluxes to Distinguish Between Metabolic Conditions

To understand how FFLs function, and therefore to explain these observed overabundances, we considered how they propagate perturbations to the input flux, *v*_1_, to the output flux, *v*_3_. We characterized this using an elasticity, or normalized gain, 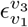, between the two fluxes for a generic FFL (Fig 3A for schematic, see Materials and Methods for model development). The magnitude and sign of this elasticity value depends on both the regulatory modes (inhibition vs activation) of the interactions comprising the loop, and the steady state effector concentrations, *x*_2_ and *x*_4_, about which the perturbation occurs. When 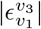 is low, *v*_3_ is weakly coupled to *v*_1_, and when it is high, *v*_3_ is strongly coupled to 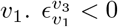 indicates that increases to *v*_1_ will cause decreases in *v*_3_ (“negative coupling”), while 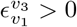 indicates that increases to *v*_1_ will cause increases in *v*_3_ (“positive coupling”).

This model demonstrates that there are metabolic states in which steady-state effector concentrations yield coupling between fluxes independent of any metabolic network structure which may connect them. SMRN FFLs thus may enable online and context-dependent control of one flux as a function of another flux, and each of the eight canonical types (Fig 3C) mediates specific flux coupling interactions across all possible effector concentrations (Fig S3). The most overabundant loop, I2, negatively couples fluxes for all effector values when 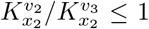 (Fig 4A, upper right panel). In this regime coupling strength is maximized when one effector concentration 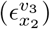 is high and the other 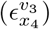 is low (Fig 4C, left panel). When 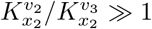, however, I2 can instead positively couple fluxes in inverse conditions (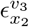 low, 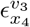 high, Fig 4A, upper left panel). This loop is unique among all FFLs in that it offers flexibility in control mode via parameter tuning (Fig S3). Depending on the parameter values, it can act as both a “brake” by reducing other fluxes or an “accelerator” by increasing them. Thus, it provides maximal flexibility for control of metabolic fluxes.

**Figure 4.**
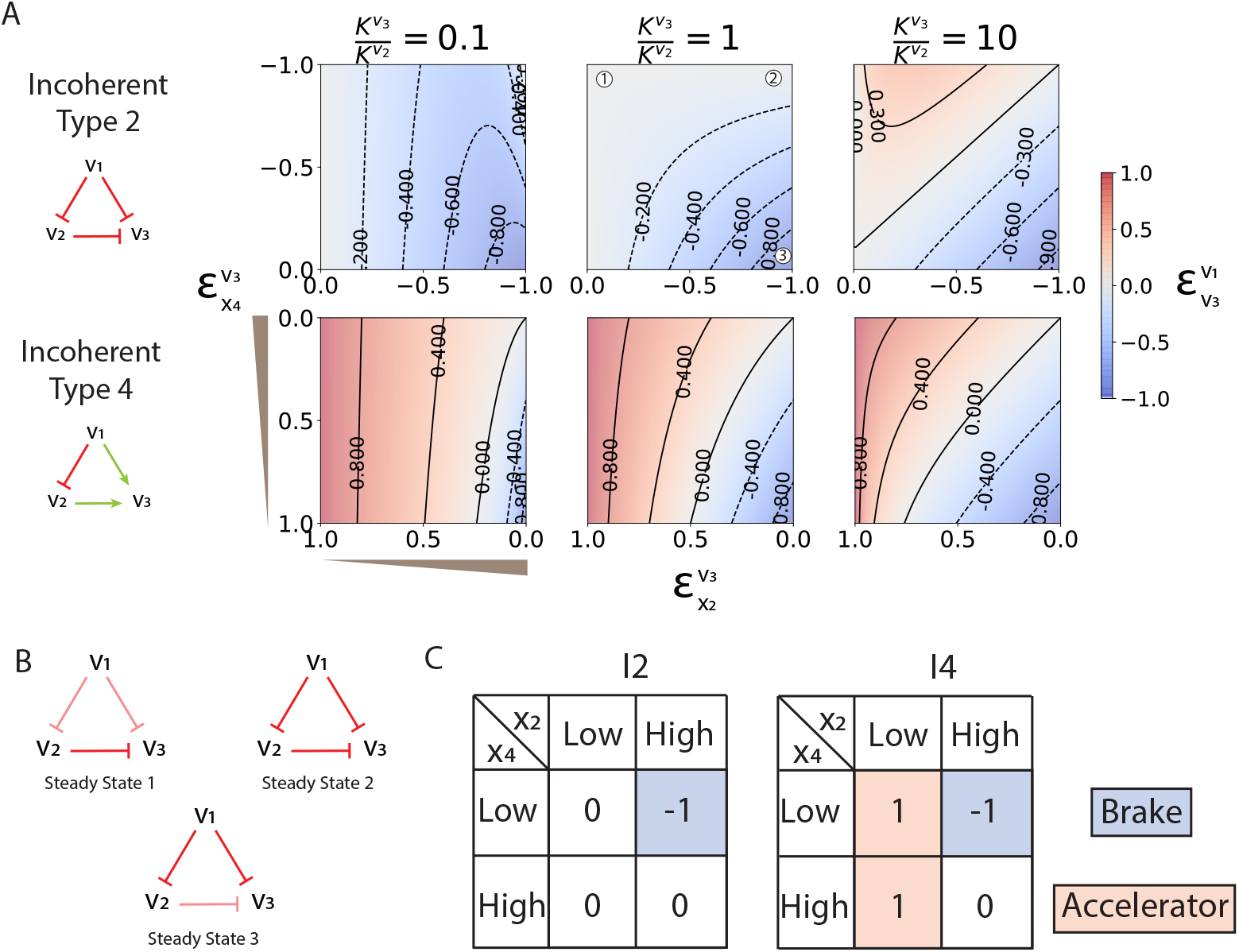
Functional properties of overabundant FFLs. (A) Flux coupling across parameter values and concentrations for I2 and I4. The relevant parameter ratio is indicated above each column in the panel grid. The x-axis of each heatmap is the elasticity value reflecting the concentration of effector 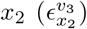 and the y-axis 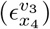 reflects the concentration of effector *x*_4_. Concentrations increase going up and the right, as indicated by the grey wedges. Coupling strength is given by the intensity of the heatmap, and sign by the colour, with blue indicating negative coupling and red indicating positive coupling. (B) Schematic representation of the three example steady-states for I2. The steady states are indicated in the middle panel of the I2 row in (A). In steady state 1, *x*_4_ has a high concentration and *x*_2_ has a low concentration. *v*_1_ is weakly coupled to *v*_3_. In steady state 2, both *x*_4_ and *x*_2_ have high concentrations. *v*_1_ is weakly coupled to *v*_3_ because the negative interactions in I2 cancel out. Finally, in steady state 3, *x*_4_ has a low concentration and *x*_2_ has a high concentration. *v*_1_ is strongly negatively coupled to *v*_3_. (C) Logic of each loop structure. Combinations of effector concentrations at given steady states yield specific coupling relationships between fluxes. I2 more often decouples fluxes, with only one effector combination yielding strong coupling. I4 more often couples fluxes, but effector concentrations tune the sign and strength of this coupling. Red boxes indicate loop function as a metabolic “accelerator”, while blue boxes indicate loop function as a metabolic “brake”. Here 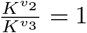.

The other overabundant loop, I4, also offers both negative and positive flux coupling, but it does so as a function of effector concentration, not parameter tuning (Fig 4B, lower panel) - it acts as a “brake” or “accelerator” depending on the current steady state effector concentrations, rather than parameter values. As one effector concentration 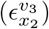 increases from zero, the I4 loop goes from mediating positive coupling to mediating negative coupling (Fig 4C, right panel). The other effector concentration 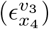 tunes the point at which this switch from positive to negative coupling occurs.

I2 and I4 offer both positive and negative coupling depending on parameter tuning and steady-state metabolite concentrations, respectively, and they are respectively the least and most robust to parameter changes of all the loops (Fig S3-5). In addition, I2 only actively couples fluxes when one steady-state effector concentration is high, and therefore most often does not couples fluxes at all. It is more often “off” than “on”, since the natural distribution of elasticity values is bimodal, with values near zero being more common than those approaching saturation [34]. Since these loops have the broadest functionality in parameter and concentration space, when they arise randomly in the SMRN they are more likely to be neutral in terms of fitness or offer a functional advantage by random chance than the other FFLs. This may explain why they are significantly overabundant in the real SMRN relative to randomized networks.

In contrast, the underrepresented loops - C2 and C4 - both only offer one mode of control across parameter space and steady-state effector concentrations (Fig S3). They act as accelerators only. Additionally, C2 amplifies flux signals, causing larger changes in *v*_3_ than those which occur in *v*_1_, regardless of parameter values (Fig S3, S4). Accelerator-only control and signal amplification could both lead to instability in the network by increasing rates in response to already-increasing rates, and/or causing extreme flux variation in response to small input perturbations. These loops are thus likely underrepresented because when they arise in the network they provide a functional disadvantage in the form of instability. The contrast between the functionality of over- and underrepresented loops highlights the fact that flexibility in coupling mode is likely a favourable feature of SMRN structures.

### Real Loops Can Distinguish Between Distinct Flux Distributions in Central Carbon Metabolism

We wanted to understand whether FFLs are relevant in real metabolic conditions, so we looked to functionalize some of these loops. We were limited to loops in central carbon metabolism because these are the only ones for which reliable parameter values are available. For each loop, we calculated relevant elasticities using steady-state metabolomic data for cells grown on 14 different carbon sources in *E. coli* [2, 8, 15]. We also made use of fluxomic data [8] which is available for 8 of these 14 carbon source to contextualize the flux coupling interactions mediated by regulatory structures and to generate hypotheses about their functions.

#### Incoherent Type 2 Loops are Tuned to Stabilize Fluxes

In the I2 loop we characterized, citrate produced by citrate synthase (CS) inhibits phosphoenolpyruvate carboxylase (PPC) and fumarase (FUM) activity, and malate produced by FUM also inhibits PPC activity (Fig 5A). This loop positively couples CS and PPC fluxes and does so as a linear function of the magnitude of CS flux (Fig 5C, *r* = 0.88, *p* = 0.004). Positive coupling between CS and PPC is necessary to balance oxaloacetate (oaa) supply with demand (Fig 5A), since assimilation into the TCA cycle of each acetyl-CoA molecule requires one oaa molecule [36]. Thus, increases to the rate at which CS generates citrate (i.e. increases to oaa demand) must be met with concomitant increases to the rate at which oaa is produced via PPC (i.e. oaa supply) to avoid instability. When oaa demand exceeds supply, indicated by an increase in CS flux, anaplerosis via PPC is stimulated by this loop to meet demand (Fig 5A, left panel). Conversely, when oaa supply exceeds demand, as indicated by a decrease in CS flux, anaplerosis is inhibited (Fig 5A, right panel).

**Figure 5.**
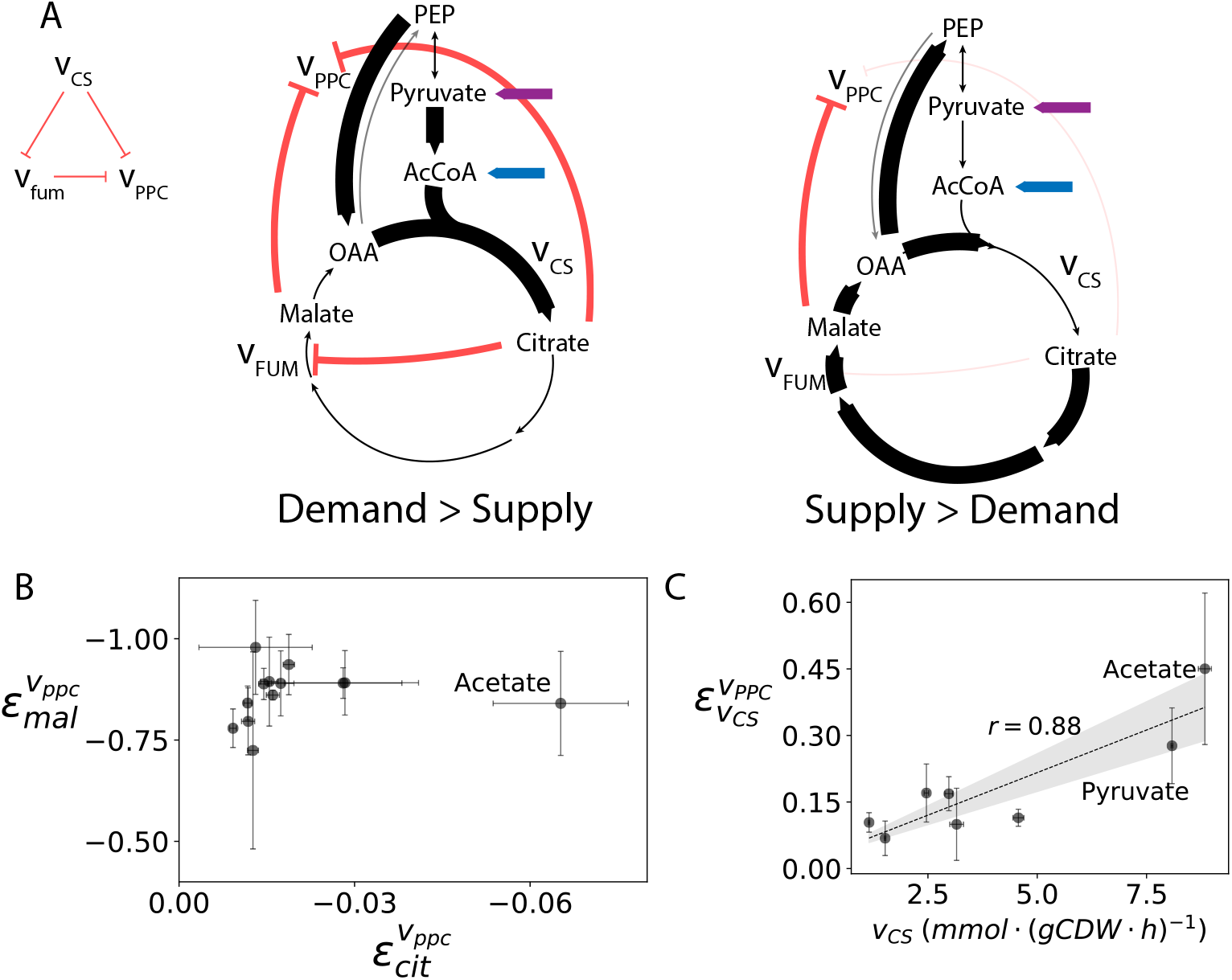
The CS-FUM-PPC loop couples oxaloacetate (oaa) supply with demand when demand is high. (A) Loop schematic and metabolic context. When influx to the TCA cycle increases, oaa demand increases, and increased anaplerosis via PPC is required to meet this demand. In contrast, when oaa supply exceeds demand, anaplerosis should be inhibited to allow for excess oaa to be used in gluconeogenic reactions. Net positive coupling of CS with PPC is required to balance supply and demand, and this loop is tuned to provide such coupling, with 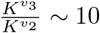 (see Fig 4). The blue arrow indicates entry into metabolism of acetate and the purple arrow indicates entry of pyrvuate, two substrates which yield high CS flux. (B) Elasticity plot for the CS-FUM-PPC loop. In all metabolic conditions, 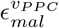 is high, so citrate concentration controls loop sensitivity. In cells fed substrates which cause citrate accumulation (e.g. acetate), coupling is maximized. (C) This leads to a linear relationship (*r* = 0.88, *p* = 0.004) between steady-state CS flux and coupling strength: when CS flux is high, coupling between CS and PPC is strongest. When CS flux is high, perturbations to this flux have the largest effect on the supply-demand balance of oaa, so maximal coupling strength is necessary.

Flux coupling via this loop is tuned to be dependent on the steady-state citrate concentration (reflected in 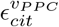), because though 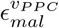 is relatively constant, indicating operation near saturation for all metabolic conditions, steady-state citrate concentration is dependent on steady-state CS flux (Fig 5B,C). As a result, flux coupling between CS and PPS is stronger as CS flux increases. As CS flux increases, the demand for oaa also increases, and marginal changes to the magnitude of CS flux have increasingly large effect on this demand, so stronger coupling is required to balance supply and demand at higher CS fluxes. As we have shown, the positive coupling essential to this system is only possible in I2 loops when 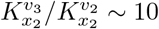, which is the case for this loop [9, 39]. Indeed, when the sensitivity of PPC to citrate inhibition is increased (K_*I*_ = 1 mM from K_*I*_ = 34.4 mM) the positive elasticity values become negative for all carbon sources, and the control landscape changes significantly, with high CS-flux states (e.g. cells fed acetate) becoming indistinguishable from other states (Fig S6). Thus this loop is well structured and tuned to function as an oaa-balancing sensor-actuator which responds to changing TCA cycle influx signalled by citrate concentration, and low-sensitivity SMR interactions are essential to this function.

#### Overlapping I4 Loops Distinguish Between Metabolic Modes

There are several I4 loops in central carbon metabolism, all of which use net pep flux (*v*_*pep*_) and net *α*-ketoglutarate (akg) flux (*v*_*akg*_) as input signals and fructose bisphosphate aldolase (FBA) as the control target (Fig 6A). We expanded our model to be able to characterize these overlapping loops (see Supplemental Material), and this revealed that they yield distinct control regimes in each of the major metabolic modes (glycolytic, gluconeogenic, glyoxylate shunt, Fig 6B). Further, a sensitivity analysis revealed that complete loops, with some low-sensitivity interactions, are required for such distinct control modes (Fig S6). This suggests that these SMRN loops are structured and tuned to use steady-state metabolite concentrations to distinguish between flux distributions in order to differentially couple fluxes based on these distributions. We found that metabolic modes are associated with either pep-dominant or akg-dominant flux coupling, suggesting that just two net rates of metabolites are required to distinguish between them.

**Figure 6.**
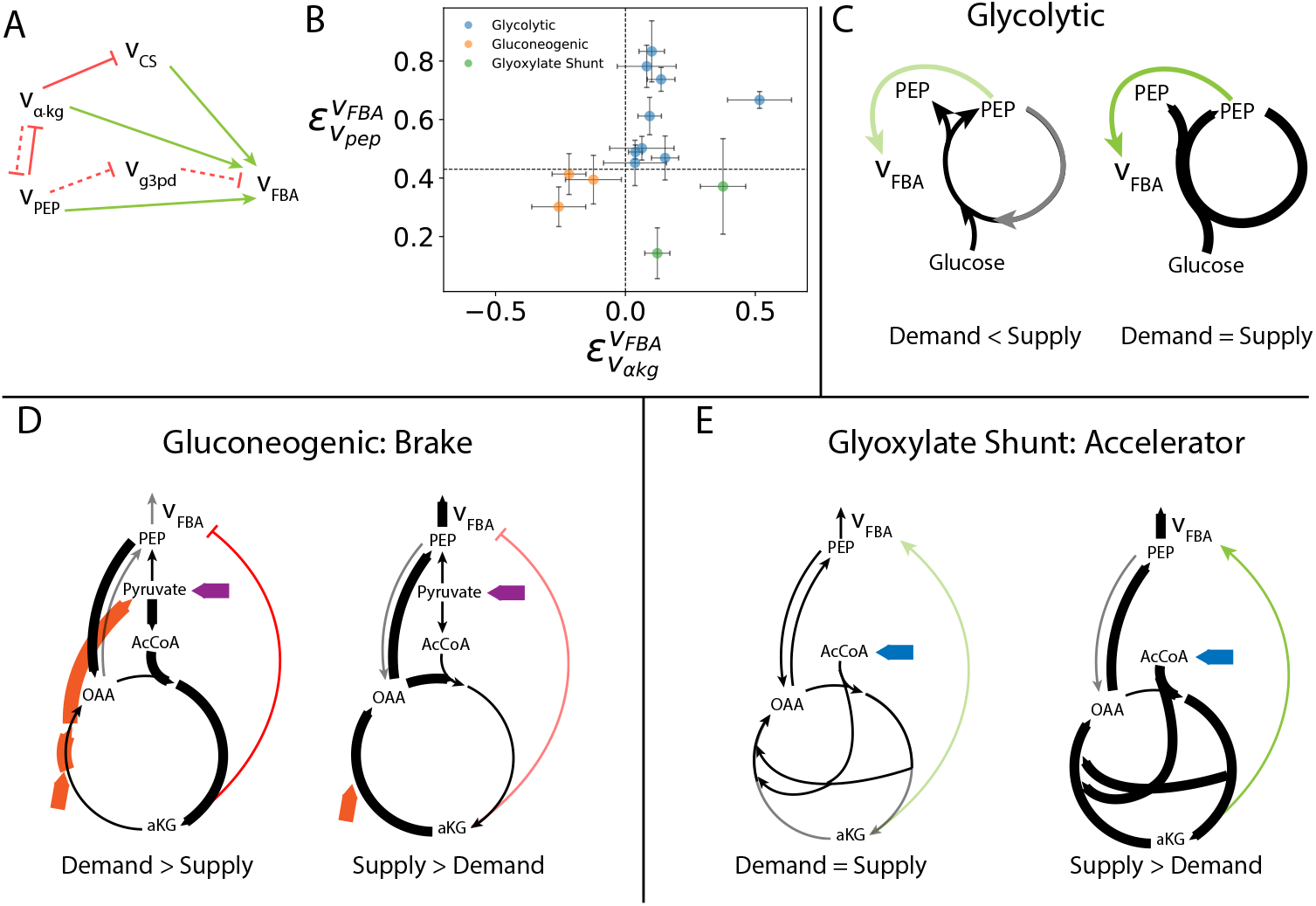
Overlapping I4 loops distinguish between metabolic modes. (A) Loop schematic. Dashed lines indicate interactions which are only relevant on some carbon sources. The *v*_*akg*_ − *v*_*pep*_ interaction is only relevant in cells fed gluconeogenic substrates and the *v*_*pep*_ − *v*_*g*3*pd*_ interaction is only relevant in cells fed glycerol. (B) These overlapping loops distinguish between major metabolic modes. In cells fed glycolytic substrates, FBA is highly sensitive to *v*_*pep*_ and weakly sensitive to *v*_*akg*_. In cells fed gluconeogenic substrates, FBA is negatively coupled to *v*_*akg*_, while in those fed substrates which use the glyoxylate shunt, FBA is positively coupled to *v*_*akg*_. Dashed lines indicate elasticity values which roughly separate metabolic modes into different control regimes. (C) *v*_*pep*_ − *v*_*FBA*_ coupling for glycolytic substrates. In these cells, pep is the substrate for an autocatalytic loop, so positive coupling of *v*_*pep*_ with *v*_*FBA*_ is an example of positive autoregulation. This may stimulate cycle flux following a new influx of glycolytic substrate (e.g. glucose) to the cycle, stabilizing glycolysis over other metabolic modes when glycolytic substrates are present. (D) Negative *v*_*akg*_ − *v*_*FBA*_ coupling in cells fed gluconeogenic substrates balances oaa supply and demand. When influx to the TCA cycle increases, oaa demand increases. *v*_*akg*_ signals this by inhibiting pep commitment to gluconeogenesis so that it can be used for anaplerosis. Conversely, when oaa supply exceeds demand, FBA activity is de-inhibted so that excess oaa can be used in gluconeogenesis. The purple and orange arrows indicate entry into metabolism of pyruvate and succinate, respectively. (E) Negative *v*_*akg*_ − *v*_*FBA*_ coupling in cells fed substrates requiring the glyoxylate shunt prevents oaa accumulation. As in the pep cycle, oaa supply and demand is balance by autocatalysis, so excess flux must be committed elsewhere to avoid accumulation. In this case, *v*_*akg*_ signals excess oaa supply that must be committed to gluconeogenesis to avoid accumulation.

Cells fed gluconeogenic substrates have akg-dominant control. In these cells, *v*_*akg*_ and *v*_*FBA*_ are negatively coupled; the *v*_*akg*_ signal is a brake. Positive *v*_*akg*_ indicates an increase in gluconeogenic substrate entering the TCA cycle in this case, and therefore an increase in oaa demand (Fig 6D, left panel). Reducing *v*_*FBA*_ activity in gluconeogenesis reduces pep demand, and may allow this increasing oaa demand to be met via anaplerosis. In contrast, negative *v*_*akg*_ indicates that akg is being consumed more quickly than it can be made, so anaplerosis is not needed and FBA activity should be increased to pull pep up through gluconeogenesis (Fig 6D, right panel). In cells using gluconeogenesis, control signalled by *v*_*akg*_ is demand-dominant, since stability is dependent on oaa demand being met.

A similar supply and demand balancing act is required in cells making use of the glyoxylate shunt (Fig 6E). While such cells also have akg-dominant control, in this case, akg is positively coupled to FBA; *v*_*akg*_ is an accelerator signal. In contrast to the gluconeogenic TCA cycle, the glyoxylate shunt is an autocatalytic cycle because it produces two oaa for every one that enters. It is therefore self-sustaining, and its stability depends on excess flux leaving the cycle [1], rather than sufficient flux supplying the cycle. In cells using the glyoxylate shunt, increasing akg flux indicates excess supply flux (Fig 6E, right panel), since these cells do not need an anaplerotic source of oaa. In this context, positve coupling of akg with FBA helps draw excess flux up through gluconeogenic reactions when it is signalled by increasing akg. When supply is not in excess, FBA is not strongly activated (Fig 6E, left panel). In cells using the glyoxylate shunt, control is supply-dominant, since excess supply must be managed. The overlapping I4 loop structure allows both supply-and demand-dominant control modes to arise from one regulatory system because of the flexibility of the I4 FFL.

In contrast to both these metabolic modes, cells fed glycolytic substrates have pep-dominant control, with akg influence on FBA near zero (with exception for cells fed glycerol). *v*_*pep*_ is strongly positively coupled to *v*_*FBA*_ in this case, so when pep accumulates, *v*_*FBA*_ increases and vice versa. Since *v*_*FBA*_ directly influences pep supply, coupling of *v*_*pep*_ to *v*_*FBA*_ is an example of positive autoregulation, and may serve any number of purposes. For example, this coupling interaction may prevent changes to *v*_*FBA*_ unless changes to *v*_*pep*_ are sustained [7]. This may prevent spurious transitions to glycolysis from other metabolic modes. However, it is difficult to assign functionality to this particular interaction without dynamic data.

As we have shown, I4 can yield both positive and negative coupling relationships and the transition between these is dependent on steady-state effector concentrations. As a result, the I4 loops in central carbon metabolism provide context-dependent control as a function of flux distributions signalled by steady-state metabolite concentrations. In all three major metabolic modes, these loops are well structured to balance supply and demand fluxes of key stability-defining metabolites, oaa and pep, by making use of signals of imbalanced supply and demand fluxes of these metabolites.

## Discussion

While it is generically understood that small-molecule regulation is necessary for the robust operation of metabolism, little has been done to uncover the fundamental mechanisms by which the SMRN achieves this. Here, we show that regulatory signals arise from fewer highly connected notes whereas the reactions being regulated are distributed across the network. With our structural approach, we have also shown that the SMRN is locally organized to achieve the goal of measuring and responding to perturbations in fluxes. This goal is consistent with both the abundance of feedforward loops we have observed in the network, and the global redistribution of node influence that we have shown defines global SMRN structure. While small-molecule regulation has canonically been described in the context of end-product feedback inhibition, our analysis complements this view by demonstrating how feedforward signalling at the network level could be useful in the sensing and control of fluxes.

In core metabolism, these loop structures appear to enable the rapid sensing of perturbations to external and internal states and coordination of short-term responses. In particular, they are able to couple relevant fluxes to perturbations in the net fluxes of key metabolites in a way that is dependent on metabolic context. These key loops act to both increase and decrease target fluxes in response such perturbations - they can act as both metabolic “accelerators” or metabolic “brakes” to maintain stable operation across metabolic modes. The metabolites involved in the loops we have characterized here have been previously identified as probable flux sensors based on their variable concentrations and/or roles as effectors [16, 21]; we have provided mechanistic evidence in support of this hypothesized function by showing how changing fluxes indicated by these metabolites could be sensed and interpreted locally by the SMRN.

More broadly, the ability for metabolic networks to sense and rapidly respond to fluxes via metabolites appears to be widespread, as we have found that more than half of all regulated reactions participate in small-molecule regulatory feedforward loops, and these all have the capability to mediate input-output flux perturbation relationships. This suggests that metabolic networks could be generally capable of sensing the direction and magnitude of internal and external flux perturbations and using this information to coordinate the dynamic response at the metabolic level.

As we have shown with the examples we characterized, small-molecule regulatory feedforward loops are strongly relevant to metabolic cycles. In such metabolic structures, stability is dependent on online balancing of rates of supply and demand, and flux coupling may be essential to ensure that this balance is achieved. As metabolic engineering is expanded into the engineering and design of cycles, consideration of this balancing act will become paramount. For example, recent investigation into synthetic carbon fixation cycles in several organisms has yielded promising results [28, 35]. However, continued optimization of these cycles – and others – will require engineered mechanisms for balancing supply and demand fluxes. We have demonstrated how SMR can be locally structured to achieve this.

Our approach is limited in a few key ways. Purely structural analyses of metabolism necessarily ignore its dynamic nature: edges and nodes are transient and context-dependent in real metabolisms as gene expression turns on and off, metabolites are depleted, etc. It is therefore difficult to extend the capabilities of SMR structures in central metabolism to the entire network without significantly more detailed understanding of each individual interaction and protein involved. Thus, we have sketched out the general design principles and systems-level function of the SMRN, not its precise operation with our analysis. Additionally, as with other network reconstructions, ours is biased by the available information describing SMR interactions. While the *E coli* network is by far the most well-validated, it is possible that the enrichments we report are biased by our current knowledge of metabolism, or that unknown interactions could complicate the mechanisms of interaction we propose here in real systems. Regardless, simplifying analyses of the type we have presented could be useful in future applications requiring accurate predictions of the dynamic responses of metabolism to environmental perturbations.

## Methods

### Network Reconstruction

We parsed information about enzyme regulators for all *E coli* entries from both BRENDA [37] and the Allosteric Database (ASD) [38], including both activators and inhibitors, and any available parameter values without regard to the listed mechanism of action. We checked entries from both databases against each other, keeping all unique interactions. Most of the ASD entries were available in the much larger BRENDA dataset. The list of interactions that resulted contained many non-native regulatory compounds, such as drugs, solvents, and heavy metals. We eliminated these by cross-referencing effectors in our list against metabolites in the iJO1366 genome-scale model [32] using their KEGG IDs. Similarly, we used the gene rules, enzyme commission numbers, and/or Uniprot accession numbers in iJO1366 to assign reactions to our list of regulatory interactions based on enzyme information available in the databases.

Since we were concerned with functional relationships, we then performed a standard loss-free metabolic network minimization with reactions and metabolites in our reduced list of regulatory interactions as protected entities [5]. This yielded a net reduction of reactions in the network from 1366 to 955, with the addition of 45 reactions from a universal list. The resulting bipartite network captured sensitivities of fluxes to perturbations, both through mass exchange via catalysis and through information exchange through regulation. A cursory examination revealed that of the ∼ 18000 regulatory edges, nearly 80% were mediated by carrier molecules such as adenosine phosphates, and NAD(P) and NAD(P)H. It is well-known that these molecules play significant regulatory roles, so we excluded all interactions they mediate. This reduced the number of regulatory interactions to 3985. The final network is thus made up of 955 reactions connected by 4400 catalytic edges, of which 623 reactions comprising the SMRN are connected by 3985 regulatory edges.

### Randomized Ensemble Generation

We used an edge-swapping method to generate a randomized ensemble of 1000 networks. In this method, two pairs of connected nodes are chosen at random and their edges are swapped provided there is no cross-connectivity between them in the first place. This swapping is performed four times for every edge in the network to ensure full coverage. To ensure that the resulting ensemble was indeed degree-preserved, we compared the in-and out-degree distributions of the ensemble networks to those of the real network using the K-S test. We counted the fraction of ensemble networks for which the null hypothesis that the distributions were from the same sample could not be rejected at a significance of 5% and report this as the significance of degree-preservation. As shown in Fig S1, 99.3% of the in-degree distributions in the ensemble could not be distinguished from those of the real network (*p* = 0.007) and the same was true for 99.5% of the out-degree distributions (*p* = 0.005).

### Topological Analysis

Topological overlap between any two nodes, *i, j*, in a network is defined as:

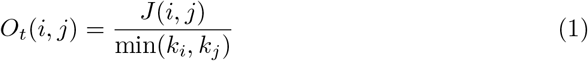

Where *J* (*i, j*) is the number of shared connections between nodes *i* and *j*, and *k*_*i*_ and *k*_*j*_ are the number of edges that nodes *i* and *j* have, respectively.

Since *O*_*t*_ is defined for undirected networks, we first converted the catalytic subnetwork to an undirected network using a built-in function in the Python Networkx package, then calculated *O*_*t*_ for each pairing of nodes in the catalytic subnetwork. Then we clustered these using the Euclidean distance to generated the topological overlap matrix (TOM) for the catalytic subnetwork. We also integrated the undirected regulatory and catalytic subnetworks by simply adding the regulatory edges to a new network containing the catalytic edges and recalculated the TOM for the integrated network to visualize the topological redistribution caused by regulatory integration with the metabolic network.

For the topological comparison of different regulatory classes, we used the average *O*_*t*_, 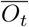, for pairings of each node with all other nodes across the entire metabolic network:

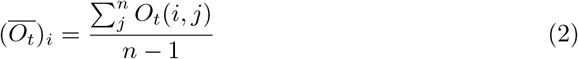

For node *i* in a network of *n* nodes with index *j*.

We also found 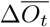 for connections between nodes by simply subtracting 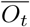 of the target of the regulatory interaction from that of the source of the interaction for every node pairing in the real SMRN:

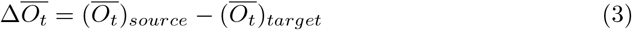

We found these differences for node pairings in each network in our randomized ensemble as well. Since it would be impossible to visualize 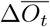 distributions for each of the randomized networks, a sample distribution is presented in Fig 2C. To ensure that the real 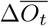 distributions were different from those of networks in the randomized ensemble, we compared 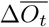 distributions for real pairings against random pairings in each of the randomized networks and calculated the significance as the number of networks in the ensemble for which the real and randomized distributions could not be distinguished. In the ensemble of 1000 randomized networks, none had 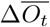 distributions with no significant difference compared to the pairings in the real network, thus *p <* 0.001.

### Flux Coupling Model

We considered the effect of a perturbation to the net input flux, *v*_1_, such that *v*_1_ → *v*_1_+*δv*_1_, on the internal flux, *v*_2_, and the output flux, *v*_3_, in a generic feedforward loop structure. Such a perturbation reflects a change in the balance of source and sink fluxes for effector *x*_2_: *δv*_1_ *>* 0 indicates a net accumulation (that need only be transient) of *x*_2_, while *δv*_1_ *<* 0 indicates decreasing *x*_2_. While most steady state concentrations are not reflective of the magnitude of their producing or consuming fluxes alone, in a transient situation with only one of these changing (due to noise, upstream flux increase, etc), the transient concentration will be reflective of the change in flux until other rates adjust due to the changes in concentrations caused by flux perturbation. Thus transient changes in the rate of accumulation of *x*_2_ in this scenario can be propagated to the other fluxes *v*_2_ and *v*_3_ via the regulatory interactions in the loop. The effect of this perturbation on the internal flux, *v*_2_, is given by the product of the sensitivity of *v*_2_ to changes in *x*_2_ and the perturbation to *x*_2_ caused by *δv*_1_:

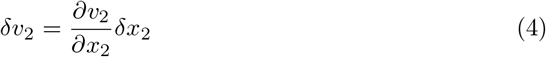

A similar sensitivity approach relates perturbations in *x*_2_ to perturbations in *v*_1_:

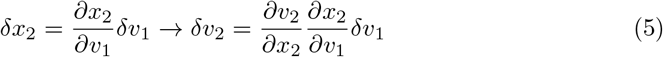

Multiplying this expression by the appropriate scaling factors yields

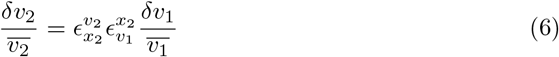

Where the overbar indicates average steady-state values, and the *ϵ* are elasticities characterizing the logarithmic gain in the superscript variable for a change about the steady state in the subscript variable:

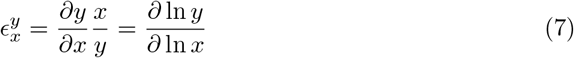

Similarly, we can examine how the perturbation will be propagated to *v*_3_, considering that this flux is sensitive to changes in both effector concentrations:

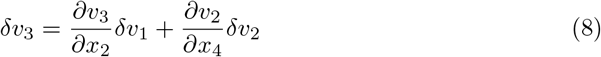

Or, in normalized form:

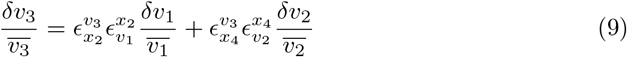

Substitution of the above expression for *δv*_2_ simplifies this equation:

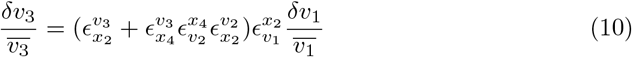

Thus, perturbations in *v*_1_ about a steady-state value will be propagated by the feedforward loop structure to the output flux under specific metabolic conditions defined by the elasticity values of the interactions it contains. In particular, the factor

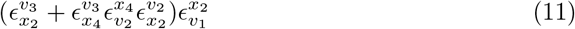

must be non-zero for perturbations to *v*_1_ to be propagated to *v*_3_ via regulatory interactions.

To simplify, we take 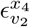 and 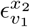 to be 1, which represents the simplified case in which a unit change in flux *v*_1_ causes a transient unit change in concentration *x*_2_. This requires assumptions of both 1:1 stoichiometry, and that *v*_1_ be the only changing flux. This yields:

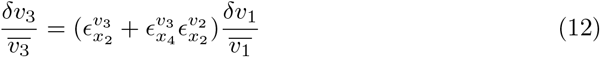

We can cast eq 12 as a new elasticity value relating fluxes by dividing both sides by 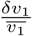 and taking the limit of *δ* → 0

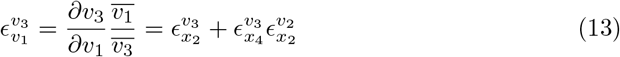

Which reveals that the normalized gain in *v*_3_ relative to a change in *v*_1_ is dependant on the product of the two elasticities in series in the feedforward loop, 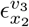 and 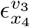, and the sum of the elasticities in parallel in the loop. These elasticities have specific forms which are dependent on regulatory mode, derived previously [34]. For inhibition by *x*_*I*_:

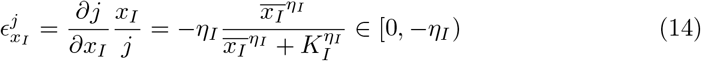

And for activation by *x*_*A*_:

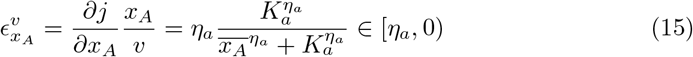

where the *η* are the Hill coefficients and the *K* are the effector binding parameters.

With these forms in place, we calculated the value of 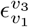 across the entire range of possible effector concentrations by noting that transforming concentrations into elasticities also transforms infinite concentration ranges into finite elasticity ranges. That is, as 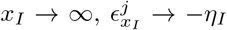 and as 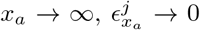. Thus, to examine how a given feedforward loop behaves, we select a value for 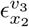 and calculate 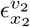 based on the loop architecture, then calculate 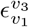 for all possible values of 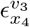. In the main text, *η* is always assumed to be 1. See Supplemental Material for a discussion of the effect of changing *η* on loop function.

### Feedforward Loop Enumeration and Identification

Since we were looking for the frequency of known FFLs in the SMRN, we did not use any motif-finding algorithms, but instead searched for loops directly by finding overlapping common neighbours for each node in the SMRN and identifying the type of regulatory interaction (activation or inhibition) connecting them. This allowed us to count unique loop-involved nodes and to count loops of each type, both in the randomized ensemble and in the real network. For the real network, this yielded 4569 feedforward loops, many of which share common input and output nodes. Of these, 94% are I2 and I4 loops.

### Feedforward Loop Functional Analysis

We used a previously collated data set comprising three reference metabolomic data sets [2,8, 15] and a reference fluxomic data set [8]. This data was used to calculate elasticities using equation 13, with the specific form of each interaction based on the interaction mode (see Supplemental Material), and *η* = 1 assumed. Since data were collated from elsewhere, we propagated error to estimate variability in the elasticities we report. This has the benefit of overestimating the variability of these values. Unfortunately most references for the regulatory parameters did not present measured variability, so we could not estimate the error associated with these values. The correlation presented in Fig 5 was performed using simple linear regression analysis in Python 3.7. All code is available as a Jupyter notebook at https://github.com/lmse/SMRN, and curated data is available as a supplemental file with this publication.

## Author contributions statement

C.E. conceived of the project, developed the computational methods, conducted the research, analyzed the data, and prepared the original draft of the manuscript and the revision. R.M. contributed to the formulation of the overall research goal, provided supervision, analyzed the data, assisted with framework development, data visualization, writing and revising the paper, acquired funding for this work. All authors reviewed the manuscript.

## Acknowledgements

Authors acknowledge funding from NSERC through the Discovery Grants program and through the M3 CREATE program. RM also acknowledges funding from Genome Canada and the Ontario government.

## References

1. U. Barenholz, D. Davidi, E. Reznik, Y. Bar-On, N. Antonovsky, E. Noor, and R. Milo. Design principles of autocatalytic cycles constrain enzyme kinetics and force low substrate saturation at flux branch points. eLife, 6, feb 2017.

2. B. D. Bennett, E. H. Kimball, M. Gao, R. Osterhout, S. J. Van Dien, and J. D. Rabinowitz. Absolute metabolite concentrations and implied enzyme active site occupancy in Escherichia coli. Nature Chemical Biology, 5(8):593–599, 2009.

3. V. Chubukov, L. Gerosa, K. Kochanowski, and U. Sauer. Coordination of microbial metabolism. Nature Reviews Microbiology, 12(5):327–340, may 2014.

4. V. Chubukov, M. Uhr, L. Le Chat, R. J. Kleijn, M. Jules, H. Link, S. Aymerich, J. Stelling, and U. Sauer. Transcriptional regulation is insufficient to explain substrate-induced flux changes in Bacillus subtilis. Molecular systems biology, 9(1):709, nov 2013.

5. P. Erdrich, R. Steuer, and S. Klamt. An algorithm for the reduction of genome-scale metabolic network models to meaningful core models. BMC Systems Biology, 9(1), 2015.

6. O. Gallego, M. J. Betts, J. Gvozdenovic-Jeremic, K. Maeda, C. Matetzki, C. Aguilar-Gurrieri, P. Beltran-Alvarez, S. Bonn, C. Fernández-Tornero, L. J. Jensen, M. Kuhn, J. Trott, V. Rybin, C. W. Müller, P. Bork, M. Kaksonen, R. B. Russell, and A.-C. Gavin. A systematic screen for protein–lipid interactions in Saccharomyces cerevisiae. Molecular Systems Biology, 6:430, nov 2010.

7. R. Gao and A. M. Stock. Overcoming the Cost of Positive Autoregulation by Accelerating the Response with a Coupled Negative Feedback. Cell Reports, 24(11):3061–3071.e6, sep 2018.

8. L. Gerosa, B. Haverkorn van Rijsewijk, D. Christodoulou, K. Kochanowski, T. Schmidt, E. Noor, and U. Sauer. Pseudo-transition Analysis Identifies the Key Regulators of Dynamic Metabolic Adaptations from Steady-State Data. Cell Systems, 1(4):270–282, oct 2015.

9. E. W. Gold and T. E. Smith. Escherichia coli phosphoenolpyruvate carboxylase: Effect of allosteric inhibitors on the kinetic parameters and sedimentation behavior. Archives of Biochemistry and Biophysics, 164(2):447–455, oct 1974.

10. S. Goyal, J. Yuan, T. Chen, J. D. Rabinowitz, and N. S. Wingreen. Achieving optimal growth through product feedback inhibition in metabolism. PLoS Computational Biology, 6(6):1–12, 2010.

11. S. Grimbs, J. Selbig, S. Bulik, H.-G. Holzhütter, and R. Steuer. The stability and robustness of metabolic states: identifying stabilizing sites in metabolic networks. Molecular systems biology, 3:146, 2007.

12. A. Gupta, I. M. Reizman, C. R. Reisch, and K. L. Prather. Dynamic regulation of metabolic flux in engineered bacteria using a pathway-independent quorum-sensing circuit. Nature Biotechnology, 35(3), 2017.

13. A. Gutteridge, M. Kanehisa, and S. Goto. Regulation of metabolic networks by small molecule metabolites. BMC Bioinformatics, 8:1–17, 2007.

14. S. R. Hackett, V. R. Zanotelli, W. Xu, J. Goya, J. O. Park, D. H. Perlman, P. A. Gibney, D. Botstein, J. D. Storey, and J. D. Rabinowitz. Systems-level analysis of mechanisms regulating yeast metabolic flux. Science, 354(6311), 2016.

15. K. Kochanowski, L. Gerosa, S. F. Brunner, D. Christodoulou, Y. V. Nikolaev, and U. Sauer. Few regulatory metabolites coordinate expression of central metabolic genes in Escherichia coli. Molecular Systems Biology, 2017.

16. K. Kochanowski, B. Volkmer, L. Gerosa, B. R. Haverkorn van Rijsewijk, A. Schmidt, and M. Heinemann. Functioning of a metabolic flux sensor in Escherichia coli. Proceedings of the National Academy of Sciences, 110(3):1130–1135, 2013.

17. R. A. Laskowski, F. Gerick, and J. M. Thornton. The structural basis of allosteric regulation in proteins. FEBS Letters, 583(11):1692–1698, 2009.

18. X. Li, T. A. Gianoulis, K. Y. Yip, M. Gerstein, and M. Snyder. Extensive in vivo metabolite-protein interactions revealed by large-scale systematic analyses. Cell, 143(4):639–650, 2010.

19. J. E. Lindsley and J. Rutter. Whence cometh the allosterome? Proc. Natl. Acad. Sci. U.S.A, 103(28):10533–10535, 2006.

20. H. Link, K. Kochanowski, and U. Sauer. Systematic identification of allosteric protein-metabolite interactions that control enzyme activity in vivo. Nature Biotechnology, 31(4):357–361, 2013.

21. A. Litsios, Á.D. Ortega, E. C. Wit, and M. Heinemann. Metabolic-flux dependent regulation of microbial physiology. Current Opinion in Microbiology, 42:71–78, apr 2018.

22. Y.-Y. Liu, J.-J. Slotine, and A.-L. Barabási. Controllability of complex networks. Nature, 473(7346):167–173, may 2011.

23. D. Machado, M. J. Herrgård, and I. Rocha. Modeling the Contribution of Allosteric Regulation for Flux Control in the Central Carbon Metabolism of E. coli. Frontiers in Bioengineering and Biotechnology, 3(October):1–11, 2015.

24. R. Mahadevan and B. O. Palsson. Properties of Metabolic Networks: Structure versus Function. Biophysical Journal, 88(1):L07–L09, jan 2005.

25. S. Mangan and U. Alon. Structure and function of the feed-forward loop network motif. Proceedings of the National Academy of Sciences of the United States of America, 100(21):11980–11985, 2003.

26. S. Maslov and K. Sneppen. Specificity and stability in topology of protein networks. Science, 296(5569):910–913, may 2002.

27. P. Millard, K. Smallbone, and P. Mendes. Metabolic regulation is sufficient for global and robust coordination of glucose uptake, catabolism, energy production and growth in Escherichia coli. PLoS Computational Biology, 13(2):1–24, 2017.

28. T. E. Miller, T. Beneyton, T. Schwander, C. Diehl, M. Girault, R. McLean, T. Chotel, P. Claus, N. S. Cortina, J.-C. Baret, and T. J. Erb. Light-powered CO2 fixation in a chloroplast mimic with natural and synthetic parts. Science, 368(6491):649–654, may 2020.

29. H. N. Motlagh, J. O. Wrabl, J. Li, and V. J. Hilser. The ensemble nature of allostery. Nature, 508(7496):331–339, 2014.

30. Y. V. Nikolaev, K. Kochanowski, H. Link, U. Sauer, and F. H-T Allain. Systematic identification of protein-metabolite interactions in complex metabolite mixtures by ligand-detected nuclear magnetic resonance spectroscopy. Biochemistry, 55:17, 2016.

31. T. Orsak, T. L. Smith, D. Eckert, J. E. Lindsley, C. R. Borges, and J. Rutter. Revealing the Allosterome: Systematic Identification of Metabolite–Protein Interactions. Biochemistry, 51(1):225–232, jan 2012.

32. J. D. Orth, T. M. Conrad, J. Na, J. A. Lerman, H. Nam, A. M. Feist, and B.Ø. Palsson. A comprehensive genome-scale reconstruction of <i>Escherichia coli</i> metabolism—2011. Molecular Systems Biology, 7(1):535, jan 2011.

33. I. Piazza, K. Kochanowski, V. Cappelletti, T. Fuhrer, E. Noor, U. Sauer, and P. Picotti. A Map of Protein-Metabolite Interactions Reveals Principles of Chemical Communication. Cell, 172:358–372, 2018.

34. E. Reznik, D. Christodoulou, J. E. Goldford, E. Briars, U. Sauer, D. Segré, and E. Noor. Genome-Scale Architecture of Small Molecule Regulatory Networks and the Fundamental Trade-Off between Regulation and Enzymatic Activity. Cell reports, 20(11):2666–2677, sep 2017.

35. A. Satanowski, B. Dronsella, E. Noor, B. Vögeli, H. He, P. Wichmann, T. J. Erb, S. N. Lindner, and A. Bar-Even. Awakening a latent carbon fixation cycle in Escherichia coli. Nature Communications 2020 11:1, 11(1):1–14, nov 2020.

36. U. Sauer and B. J. Eikmanns. The PEP—pyruvate—oxaloacetate node as the switch point for carbon flux distribution in bacteria. FEMS Microbiology Reviews, 29(4):765–794, sep 2005.

37. I. Schomburg, L. Jeske, M. Ulbrich, S. Placzek, A. Chang, and D. Schomburg. The BRENDA enzyme information system–From a database to an expert system, nov 2017.

38. Q. Shen, G. Wang, S. Li, X. Liu, S. Lu, Z. Chen, K. Song, J. Yan, L. Geng, Z. Huang, W. Huang, G. Chen, and J. Zhang. ASD v3.0: unraveling allosteric regulation with structural mechanisms and biological networks. Nucleic Acids Research, 44(D1):D527–D535, jan 2016.

39. J. W Teipel and R. L. Hilt. The Number of Substrate-and Inhibitor-binding Sites of Fumarase*. Journal of Biological Chemistry, 243(21):6679–6683, 1968.

